# Fast, robust and precise 3D localization for arbitrary point spread functions

**DOI:** 10.1101/172643

**Authors:** Yiming Li, Markus Mund, Philipp Hoess, Ulf Matti, Bianca Nijmeijer, Vilma Jimenez Sabinina, Jan Ellenberg, Ingmar Schoen, Jonas Ries

## Abstract

We present a fitter for 3D single-molecule localization of arbitrary, experimental point spread functions (PSFs) that reaches minimum uncertainty for EMCCD and sCMOS cameras, and achieves more than 10^5^ fits/s. We provide tools to robustly model experimental PSFs and correct for depth induced aberrations, which allowed us to achieve an unprecedented 3D resolution with engineered astigmatic PSFs, and acquire high quality 3D superresolution images even on standard microscopes without 3D optics.

---

As most biological structures have a three-dimensional (3D) organization, their comprehensive investigation by superresolution microscopy requires not only a high lateral, but also a high axial resolution. Therefore, several methods have been developed to extend single molecule localization microscopy (SMLM) to 3D. Most approaches extract the z-position from the shape of an engineered point spread function (PSF)^1–3^. However, these methods are prone to produce artifacts, which are often a consequence of a mismatch between the theoretical PSF model and the experimental PSF. Thus, an accurate PSF model is a prerequisite for accurate 3D localization of single molecules. Any analytical PSF model is an approximation at some level, even with a (computationally expensive) consideration of optics and sample induced aberrations. Therefore, approaches using an experimentally acquired PSF for single molecule localization have been developed, such as PSF correlation^4^, phase retrieval^5,6^, or interpolated PSFs^7–11^. The latter approach is especially promising, as it can model any experimental or theoretical PSF. However, at the moment, all these methods are either of limited accuracy, lack robustness or are rather slow, and the generation of an accurate PSF model is challenging. Thus, in contrast to simple Gaussian PSF approximations, experimental PSFs are rarely used for SMLM.

Experimental PSF models can be generated from a z-stack of beads of high signal to noise ratio. This is not trivial, as any imperfections easily lead to artifacts, often seen as deformations and stripes in the reconstructed images (**Supplementary Fig. 1)**. Thus, we developed a simple tool to generate a robust and accurate PSF model from several bead stacks across different field of views, which avoids those artifacts by proper averaging and regularization (**Supplementary Fig. 1, Online Methods**).

Most importantly, we implemented a robust fitting algorithm for cubic spline (cspline) interpolated PSF models. It reaches ultra-fast computational speeds by implementation on the graphics processing unit (GPU) (**Fig. 1 a**) and achieves the theoretical limit in localization precision, the Cramér-Rao lower bound (CRLB, **Supplementary Fig. 2**). Compared to a Gaussian PSF model, the spline-interpolated PSF model improved both the localization accuracy, as well as the localization precision, on simulated (**Supplementary Fig. 3**) and experimental (**Supplementary Fig. 4**) data. In addition, it allows for an accurate and precise determination of the number of photons per localization, which cannot be accurately recovered using a Gaussian PSF approximation (**Supplementary Fig. 5**). This is an important improvement for any kind of ratiometric superresolution imaging, where photon numbers in two channels are used to extract e.g. color^12^, polarization anisotropy^13^ or *z*-positions of single molecules^14^.

**Figure 1.**
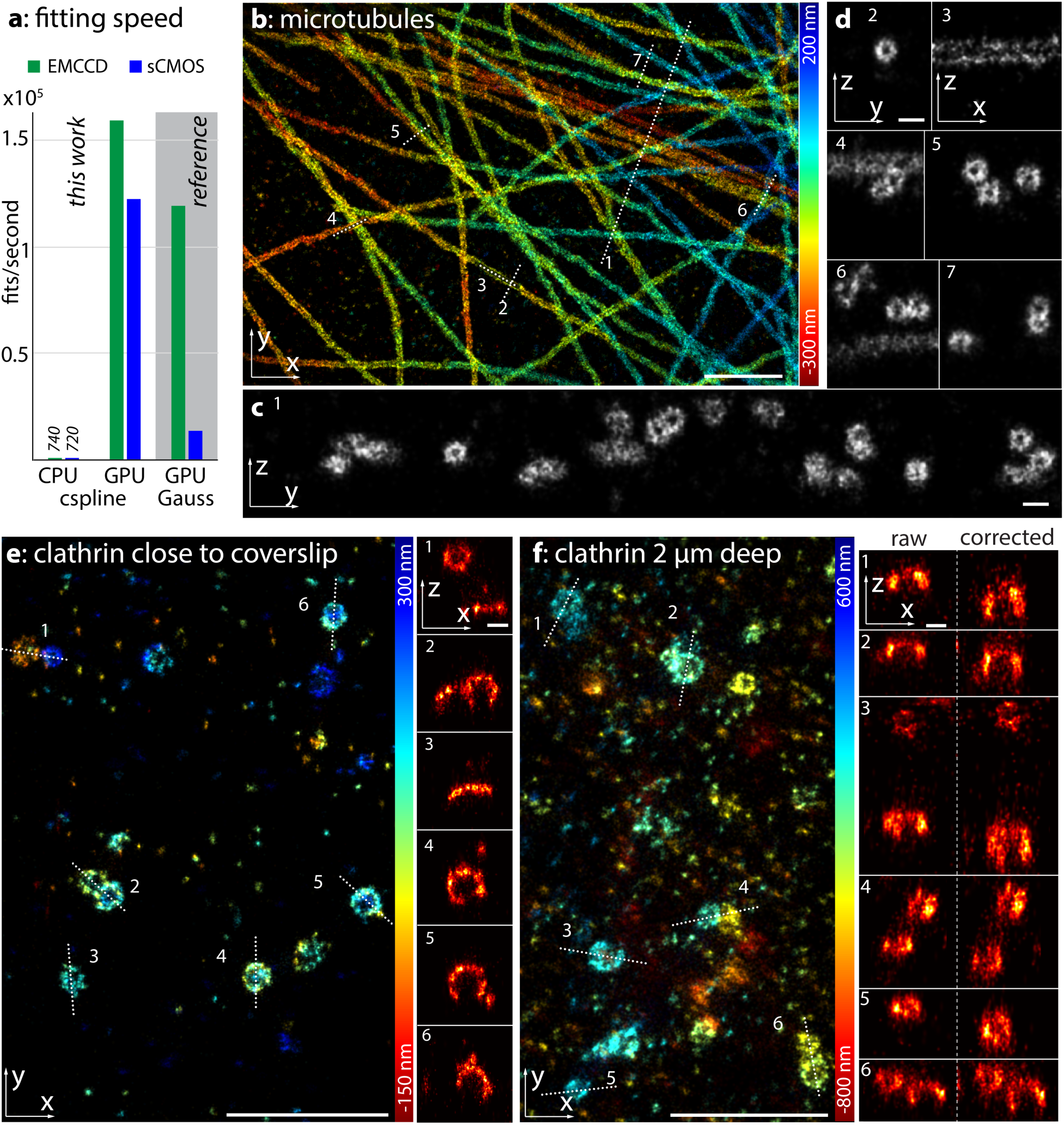
**(a)** Performance of our L-M implementation of a single-molecule fitter using cspline-interpolated experimental PSFs in comparison to a Newton implementation of a Gaussian PSF model for EMCCD^15^ and sCMOS^17^ cameras. Fits/s were measured on a i7-5930 CPU and a GTX1070 consumer graphics card. **(b)** Microtubules, labeled with α-and β-tubulin primary, and DNA-coupled secondary antibodies, and imaged using the DNA-PAINT^22^ approach. The localizations are color coded according to their z-positions. Corresponding localization precisions and profiles can be found in **Supplementary Fig. 11. (c, d)** side-view reconstructions along the lines denoted in **(b)** clearly reveal the hollow, cylinder-shaped structure of the immunolabeled microtubules. **(e)** Clathrin-coated pits, close to the coverslip, immunolabeled with Alexa Fluor 647 conjugated antibodies and measured using the dSTORM method^23^. **(f)** Clathrin-coated pits on the upper cell membrane, imaged 2 µm above the coverslip using an oil objective, show deformations in the side-view reconstructions. After correction of aberration-induced artifacts the spherical shape of the pits is recovered. Width of the line profiles: 150 nm (**c** 1, **d** 4, 6), 200 nm (**d** 2, 5, 7), 30 nm (**d** 3), 50 nm (**e, f**). Scale bars: 1 µm (**b, e, f**) and 100 nm (**c, d**, and x-z reconstructions in **e** and **f**).

The implementation of the fitting algorithm is based on maximum likelihood estimation (MLE) for the Gaussian PSF model^15^, which we extended to cspline-interpolated PSFs. For legacy, we also incorporated the standard Gaussian PSF model. Instead of the Newton method, we re-implemented the iterative procedure with the Levenberg-Marquardt (L-M) algorithm^16^ (**Online Methods)**, which converges more rapidly (**Supplementary Fig. 6**) and performs more robustly (**Supplementary Fig. 7**). Furthermore, we included the sCMOS noise model^17^ in our MLE fitter (**Online Methods**), thus enabling the use of increasingly popular sCMOS cameras for any PSF-engineering approach in 3D SMLM. Using simulated data, we could demonstrate that a fitting bias introduced by pixel-dependent readout noise was faithfully avoided (**Supplementary Fig. 8**).

Next, we evaluated the speed of our implementation (**Fig. 1a**). The central processing unit (CPU) implementation reaches about 700 fits/s on a standard PC for a 13 × 13 fitting window. It is 3 – 30 fold faster than previous fitters for experimental PSFs^9,11^ and can be used as a backup on computers without a CUDA-enabled GPU. Our GPU implementation is about 100 times faster than the CPU implementation for all ROI sizes (**Supplementary Fig. 9**), and reaches 1.6 × 10^5^ fits/s on a consumer graphics card (**Fig. 1**). We further compared the cspline fit with the conventional elliptical Gaussian fit where the asymmetry of the Gaussian PSF is used to calculate *z*. Our cspline fit is faster than the previous fitter for a Gaussian PSF model for the EMCCD camera (**Fig. 1a**). By proper memory optimization (**Online Methods**), the correction of the readout noise for sCMOS data only decreases the fitting speed by less than 15% and it is about one order of magnitude faster than the previous GPU implementation accounting for the sCMOS noise model^17^ (**Fig. 1a** and **Supplementary Fig. 9**).

We next validated our cspline fitter on biological data using an experimental PSF model generated with the above-mentioned tool. We were able to easily resolve the hollow cylinder of immunolabeled microtubules with DNA-PAINT (**Fig. 1b-d**) and dSTORM (**Supplementary Fig. 10**). Additionally, we could visualize the precise 3D organization of clathrin-coated pits (**Fig. 1e**). Their spherical geometry makes them useful calibration standards to verify the accuracy of the *z*-calibration. To our knowledge, the achieved image quality and 3D resolution is unprecedented for PSF engineered 3D SMLM (**Fig. 1, Supplementary Fig. 10** and **11**) and only reached with a much more complex interference-based 4 Pi approach^18^.

Most researchers use oil objectives for SMLM, because of their high collection efficiency, compatibility with total internal refraction excitation and simple implementation of focus stabilization. However, the refractive index mismatch between the glass and the sample leads to strong, mostly spherical aberrations when imaging above the coverslip inside the sample. These aberrations result in systematic errors in the z-localization, which can be substantial even for a moderate imaging depth of a few micrometers (**Fig. 1f**). We developed a simple software tool to correct for these systematic errors. Our approach is to take many z-stacks of beads immobilized in a layer of gel above the coverslip and fit them with a PSF model of choice to obtain the fitted z-position in dependence on the objective position (**Online Methods, Supplementary Fig. 12**). In addition, we extract the absolute z-position of the beads above the coverslip by determining the slice in which the fitted z-position is zero. This allows us to calculate the difference between the true and the fitted z-position for many objective positions which we use to correct the measured z-positions in other measurements that use the same PSF model (**Online Methods**). Using this approach, we could correct for aberration induced fitting errors at a depth of 2 µm and fully recover the spherical geometry of clathrin-coated pits (**Fig. 1f**).

Most SMLM is performed using a standard microscope in a 2D configuration without any PSF modification. Here, a Gaussian PSF model is often employed and the fitted width of the Gaussian is used as an estimate of the *z*-position to filter out out-of-focus localizations. Recently, photometry was used to extract a measure for the fluorophores’ *z*-positions^19^, showing that even simple 2D unmodified PSFs contain abundant information on the *z*-position of the fluorophore. However, the resolution at the focus has been very low and the symmetry of the unmodified PSF prevented distinguishing fluorophores above and below the focus. Thus, the focus had to be placed below the sample, restricting the measurement to the vicinity of the coverslip. In reality, an experimental unmodified PSF is not completely symmetric, but shows subtle differences between the upper and lower half, which can break the degeneracy. Here, we reasoned that by fitting with experimentally derived PSF models, we can exploit these differences to assign a fluorophore to the correct half of the PSF. To this end, we fit every single molecule image two times, once with a *z* starting parameter above the focus and a second time with a *z* starting parameter below the focus, and select the solution with the maximum likelihood. We found that the *z*-resolution for unmodified PSFs is comparable to that of astigmatic PSFs (**Fig. 2a**). Only close to the focal plane it is decreased, albeit much less than calculated from a theoretical PSF model^20^. 5% of misassignments lead to a faint mirror image (**Fig. 2d**), which for most practical applications is negligible. When we imaged the protein Nup107 in the nuclear pore complex, we could easily resolve the nucleoplasmic and cytoplasmic rings, which are axially spaced apart by only 53 nm^21^ (**Fig. 2b-d**). Thus, our new fitter enables high-resolution 3D imaging directly on standard microscopes without any 3D optics.

**Figure 2.**
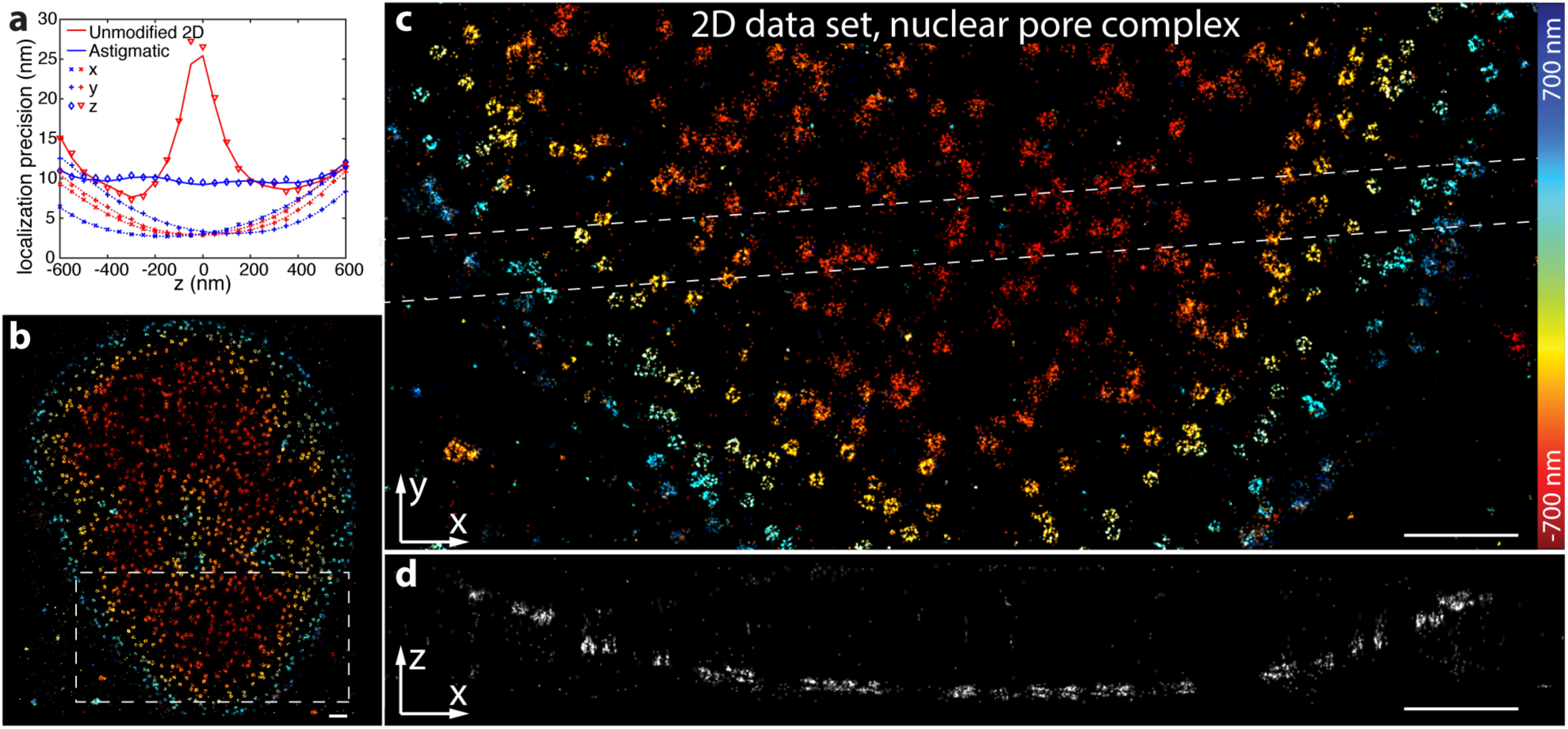
The cspline fit extracts accurate 3D positions from a simple 2D dataset with an unmodified PSF. **(a)** Localization precision in x, y and z for an astigmatic 3D and an unmodified PSF. Calculated on simulated data with 5000 photons/localization and 10 background photons/pixel. Lines represent the corresponding CRLB. (**b**) dSTORM of Nup107-SNAP labeled with BG-AF647, imaged with a standard microscope without 3D optics, overview image. (**c**) Top view reconstruction of the region denoted in (b). (**d**) Side view reconstruction of the region denoted in (c). The nucleoplasmic and cytoplasmic rings of the nuclear pore complex, spaced 53 nm apart, can be easily resolved. Corresponding localization precisions and profiles can be found in **Supplementary Fig. 11.** Scale bars: 1 µm.

To summarize, we presented a robust single-molecule fitter for arbitrary PSF models using MLE with a noise model for both EMCCD and sCMOS cameras. This allowed us to achieve an unprecedented 3D resolution and image quality using engineered astigmatic PSFs or unmodified PSFs from a standard microscope. An optimized GPU implementation achieved a fitting rate of >10^5^ fits/s, which is two orders of magnitude faster than a single threaded CPU implementation. As deformations of the PSF are included in the experimental PSF model, our fitter is robust with respect to aberrations, leading to a high accuracy even for objectives with a mediocre PSF or imperfect alignment of the microscope. Using a novel approach to correct for depth-induced aberrations, we could retain a high 3D resolution even several micrometers above the coverslip. The presented framework is not restricted to bead-stack based PSFs, but can be used in the same way to obtain and fit a spline-interpolation of an arbitrary analytical or phase retrieved PSF model, which can directly take into account aberrations.

We provide our CPU based C-code and the GPU based CUDA-code as open-source to the community (github.com/jries/fit3Dcspline.git), which can be easily incorporated in any programming language, and thus will greatly improve speed and accuracy of any single-molecule fitting software.

## ACKNOWLEDGEMENTS

We thank Johanna Mehl for helping with the sample preparation and data acquisition. This work was supported by an ERC consolidator grant (J.R.), by the EMBL Interdisciplinary Postdoc Programme (EIPOD) under Marie Curie Actions COFUND (Y.L.), the 4D Nucleome / 4DN NIH Common Fund (U01 EB021223) (J.E., J.R.) and by the European Molecular Biology Laboratory (Y.L., M.M., P.H., U.M., B.N., V.J.S., J.E., J.R.).

## AUTHOR CONTRIBUTIONS

Y.L. and J.R. conceived the approach, developed the methods, wrote the software and analyzed the data. M.M. performed the imaging of clathrin coated pits and acquired the bead images in gel. P. H. performed the imaging of nuclear pore complex. U. M. performed the DNA PAINT imaging of microtubules. B.N, V.J.S. and J.E. contributed the Nup107 cell line. I.S. contributed the custom-made DNA-PAINT antibodies. Y.L. and J. R. wrote the manuscript with the input from all other authors.

## ONLINE METHODS

### Robust averaging of experimental bead stacks

Stacks of beads, immobilized on a coverslip, were acquired in a range of ±1000 nm with respect to the coverslip. A spacing in *z* between 10 nm and 50 nm works well. Beads in each stack were segmented in a maximum intensity projected image by maximum finding and thresholding. Sub regions around each bead location were cropped. Next, we aligned each bead stack in 3D using an approach similar to single-particle averaging. To this end, the first bead is registered to the average of all bead stacks, the second bead is registered to the first bead and all remaining beads are registered to the average of the previously registered beads with sub-pixel accuracy by 3D cross-correlation of the central part of the stacks. We scaled up the central part of the cross-correlation by a factor of 20 by cubic spline interpolation and determined the x, y, and z shifts from the position of the maximum^24^. The bead stacks were shifted using cubic spline interpolation. Iteratively, bead stacks which showed a large dissimilarity from the average were identified based on the maximum value of the cross-correlation and the mean square error and excluded from the average. To eliminate the background, the minimum value of the bead stack was subtracted and the amplitude was normalized by the total (summed up) intensity of the central slice. We further regularized the bead stack by smoothing it in the z-direction with a smoothing B-spline^25^.

For astigmatic PSFs, we alternatively provide an option to register the beads in *z* based on the elliptical Gaussian fit. To this end, we determined for each bead the z-position where *σ*_*x*_(*z*) and *σ*_*y*_(*z*) of the PSF are equal. This value was used for *z*-registration and *x* and *y* registration were performed as described above with a 2D cross-correlation.

### Calculation of cspline-interpolated PSFs

Spline functions are piecewise polynomials for which high order derivatives are continuous at the knots, where the pieces connect. Cubic splines are the most commonly used splines, e.g. in computer graphics, geometric modeling, etc. Recently, this type of approximation theory has also been used for single molecule localization^9–11^. We implemented the cspline interpolation both in terms of cubic splines and cubic B-splines. A B-spline interpolation is generally less memory intensive since only one B-spline coefficient is needed in each spline interval. In comparison, (*d + 1*)*^n^* coefficients are required in each spline interval for spline polynomials, where *d* is the spline degree and *n* is the dimension. However, our implementation of a 3D fit based on cubic splines is about 2.5 times faster than the cubic B-spline form due to the fact that cubic splines are more explicit and less calculations are needed to calculate spline values and derivatives. Therefore, the software used in this work is based on cubic splines with 64 coefficients in each voxel of the 3D PSF stack.

Similar to Ref. 11, the 3D PSF is described by a three dimensional cubic spline for voxel (*i, j, k*) as follows:

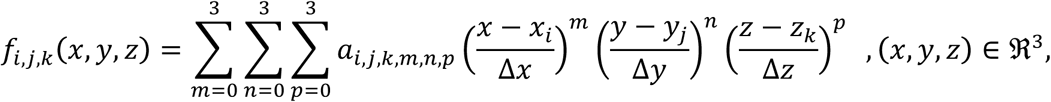

where Δ*x* and Δ*y* is the pixel size of the PSF in the object space in *x* and *y* directions, respectively. Δ*z* is the step size in the objective space in *z* direction. *x*_*i*,_ *y*_*j*_ and *z*_*k*_ are the start positions of voxel (*i, j, k*) in *x*, *y* and *z* directions, respectively.

In order to calculate the cspline coefficients, the 3D PSF stack was firstly built by averaging the bead stacks from different fields of view by 3D cross correlation and by regularization, as described above. The spline coefficients were built based on the averaged and smoothed 3D PSF stack. As 64 cspline coefficients are required to describe each voxel, we up sampled (cubic spline interpolation) each voxel 3 times in *x*, *y* and *z* directions, respectively. The 64 up sampled coordinates (including boundary of neighboring voxels) were used to calculate the 64 cspline coefficients.

### *z*-calibration of astigmatic Gaussian PSF models

Our PSF calibration tool also allows extracting *z*-positions using two widely used algorithms: a) calculate the *z*-positions directly from the calibrated *σ*_*x*_(*z*) and *σ*_*y*_(*z*) returned by the elliptical Gaussian fit; b) determine the z-positions by directly fitting the single molecules with the calibrated astigmatic Gaussian PSF model^26^.

For both calibrations, the bead stacks are fitted with an elliptical Gaussian PSF model and shifted in *z* according to their true *z*-positions where *σ*_*x*_ *z* == *σ*_*y*_(*z*). The outliers are removed based on the root mean error of *σ*_*x*_(*z*) and *σ*_*y*_(*z*) with respect to the average curves.

For algorithm a), we calculate *dσ*^2^(*z*) = *σ*_*x*_ *z* ^2^-*σ*_*y*_ *z* ^2^ and interpolate the functional relationship *z*(*dσ*^2^) by a smoothing cubic B-spline. This B-spline interpolation is then used to directly read out *z* from *d*σ^2^.

For algorithm b), *σ*_*x*_(*z*) and *σ*_*y*_(*z*) are fitted with a polynomial approximation for the astigmatic Gaussian model:

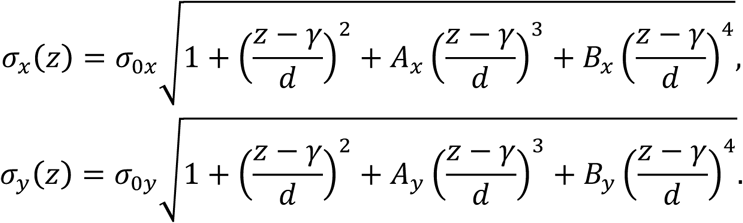

The parameters *σ*_0*x*_, *A*_x_, *B*_x_, *σ*_0*y*_, *A*_y_, *B*_y_, γ and *d* are input parameters for the Gaussian fitter, which directly returns the *z*-coordinates of the fluorophores. We follow the formula in Ref. 15 to calculate the derivatives of the parameters. However, the iterative process was re-implemented using the L-M algorithm.

## Newton and Levenberg-Marquat iterative schemes for MLE

Maximum likelihood estimation is the method of choice for fitting data with Poisson statistics^27^. The objective function for MLE is given by^16^:

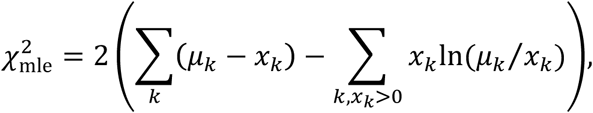

where *µ*_6_ is the expected number of photons in pixel *k* from the model PSF function, *x*_6_ is the measured number of photons. By minimizing 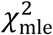, we obtain the maximum likelihood for the Poisson process.

Methods for nonlinear optimization are usually iterative. For Newton iterative schemes, the search direction *Δθ*_*i*_ of each iteration is given by^15^

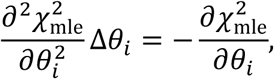

where *θ*_*i*_ is the *i*-th free fit parameter. However, computing the second derivatives is often quite difficult and can be destabilizing when the model fits badly or outlier points are contaminated that are unlikely to be compensated^28^.

An alternative method is the L-M algorithm. The L-M algorithm is often used for least square fitting as it is quick and robust. With relatively simple modifications^16^, the L-M algorithm has also been used to minimize 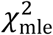. In the L-M algorithm, the second derivatives term are neglected and only the first derivatives are used. In the L-M algorithm, the update *Δθ*_*i*_ is given by

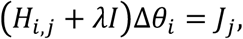

where, *H*_*i,j*_ is the Hessian matrix without the second partial derivatives term, defined as *H*_*i,j*_ = 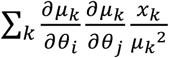, *J*_*j*_ is the Jacobian matrix defined as *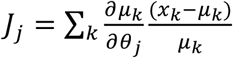*, *λ* is the damping factor, *I* is a diagonal matrix equal to the diagonal elements of the Hessian matrix. This method is more robust since a damping factor is introduced and the second derivatives do not contribute. This damping factor is increased (multiplied by 10 in this work) if an iteration step does not decrease 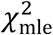 or *H*_*i,j*_ is not positive definite.

### GPU Implementation

This GPU implementation of the iterative method follows the framework developed for fitting a Gaussian PSF model using a GPU^15^. Unlike previous work for EMCCD and sCMOS noise models^15,17^, where the shared memory was used to store the molecule candidate data and readout noise map, we kept the data in the GPU global memory. Each thread is pointed to each molecule candidate and performs all the computations for each molecule candidate. No thread synchronization is required. 64 threads per block were used. The overall speed is about 1.2 times (small window size) to 10.6 times (large window size) faster (**Supplementary Fig. 9**) than for the original code where shared memory was employed for the sCMOS noise model. We assume that this is due to the compiler optimization where more registers are used and the time for copying data from the global to the shared memory is saved. Both, the CPU based C-code and the GPU based CUDA-code were compiled using Microsoft Visual Studio 2010. The software was called via Matlab (Mathworks) mex files. It was run on a personal computer using an Intel(R) Core(TM) i7-5930 processor clocked at 3.50 GHz with 64 GB memory. An NVIDIA GeForce GTX 1070 graphics card with 8.0 GB memory was used for GPU based computation.

### MLE fit using an sCMOS camera noise model

sCMOS cameras have become more and more attractive for localization microscopy due to their fast data acquisition even for large fields of view, low readout noise and relatively low price. However, their intrinsic pixel-dependent gain, offset and readout noise can create a localization bias, which has to be corrected when localizing the single molecule^17^. Gain *g*_*k*_ and offset *o*_*k*_ in pixel *k* can be taken into account when converting the camera image *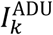* in analog digital units (ADU) into photons:

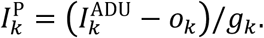

The readout noise, however, has to be taken into account during the fitting in the noise model and can be calculated from many dark camera images as the pixel-wise variance. Here, we use the same model as proposed by Huang, et al.^17^ which approximates the normal distributed readout noise (*var*_*k*_, in units of photoelectrons) with a Poisson distribution. By adding a pixel-dependent constant, *var*_*k*_, to the measured photoelectrons, one can expect the new value to approximate a Poisson distribution with a mean of *u*_*k*_ + *var*_*k*_. Here, *u*_*k*_ is the expected photon number in pixel *k* of the PSF model function. Therefore, in comparison to the conventional MLE fit for EMCCD data, only one more parameter, *var*_*k*_, is required for sCMOS data. Also *var*_*k*_ is only kept in the global memory of the GPU. Compared to the EMCCD noise model, the speed performance of the algorithm was only reduced by less than 15% by additionally accounting for the pixel dependent readout noise (**Supplementary Fig. 9**).

### Correction of depth-induced localization errors

Local squeezing or expansion in *z* are often observed in the reconstructed images when imaging deep in the cell. A common cause are depth-induced aberrations in conjunction with a bead calibration on a coverslip, which lead to a mismatch between PSF model and real PSF. However, it is a cumbersome procedure to experimentally measure the depth-dependent PSF, e.g. using an optical trap^29^. Therefore, we adopted a different strategy in which we use the imperfect PSF model for fitting, determine the fitting errors and correct for them in a post-processing step.

To calibrate the magnitude of *z*-localization errors for a specific PSF model in dependence on the imaging depth, we embedded fluorescent beads in a layer of agarose gel above the coverslip and acquired *z*-stacks at many sample positions in a range from 1 µm below to 3 µm above the coverslip with a spacing of typically 20 nm to 50 nm. Next, we determined the nominal *z*-position of each bead in the stack from the coverslip. As an estimate for the true *z*-position of a bead, we chose the frame in the stack for which the fitter returned a *z*-position of zero. By interpolation, we could achieve this with an accuracy better than the distance of the calibration planes. This measure for the *z*-position of a bead is certainly a good choice for astigmatism based 3D SMLM: as depth-induced aberrations are mostly symmetric, they do not change the asymmetry of the PSF dramatically. Thus, the focal plane, in which a bead appears symmetric is largely independent on the aberrations and thus can be defined as its nominal position. For more complex PSFs, this choice is still a good approximation. The position of the coverslip was determined from the positions of the lowest beads.

Each frame in the stack corresponds to a different focal plane. With the nominal *z*-position obtained above, we can determine the *z*-position of each bead at each frame (focal plane) and compared it with the fitted *z*-position. As we take into account many bead stacks, we can determine this *z*-correction for many combinations of fitted *z*-positions and focal plane positions (**Supplementary Fig. 12b**). We performed a robust interpolation of these data with a smoothed cubic B-spline by iteratively removing outliers with a too large distance from the smoothed surface, until no outliers were present any longer.

To correct fitted *z*-positions in an SMLM experiment, one needs to determine the approximate focal position above the coverslip. This can be achieved for instance by focusing first on dyes unspecifically bound to the glass or residual fluorescence on the glass surface, before focusing to the area of interest and reading out the difference in objective depth. Once we know the objective depth, the interpolated *z*-correction function directly returns the *z-*correction from the focal plane position and the fitted *z*-values.

**Supplementary Fig. 12c** shows a validation of this approach on beads. The left panel shows the fitted *z*-positions in dependence on their nominal distance from the focal plane, and for many beads these are not equal (root mean square (rms) error 148 nm, Pearson correlation coefficient c=0.9848). The right panel shows the corrected *z*-positions, which now show a very high correlation with the true relative positions (rms error 18 nm, Pearson correlation coefficient c=0.9993).

This correction for depth-dependent axial distortions complements previous work correcting for depth-dependent lateral distortions^30^. However, when using an experimental PSF, such lateral corrections are usually not required, as the PSF model takes into account any asymmetry.

### 3D Fitting of SMLM data acquired with standard microscopes without 3D optics

As our code includes fitting with arbitrary PSFs, it is directly applicable on 2D data, acquired with an unmodified PSF in a standard microscope. A model for the unmodified PSF can be calculated directly from bead stacks, in an anologous way to engineered PSFs. However, the unmodified PSF has a high symmetry with respect to the focal plane (**Supplementary Figure 13**), making it difficult for an iterative fitting procedure to converge through the focal plane. To overcome this problem, we fit every localization twice: once with a starting parameter for the *z*-position 500 nm above the focal plane, and once with a starting parameter 500 nm below the focal plane. The maximum likelihood is then used to select the better fit. As a real PSF is not completely symmetric^31^, this breaks the degeneracy previously encountered when extracting *z*-positions in 2D data sets from only a single photometry or PSF size parameter^19^.

Due to the rather large size of the calibration bead (100 nm) and small inaccuracies during the averaging of many bead stacks, the cspline PSF model is slightly blurred compared to a single-molecule PSF. This had no apparent effect on 3D data, but in 2D data it lead to an accumulation of fitted localizations at the focal plane. To overcome this problem, we filtered the raw images with a Gaussian kernel (standard deviation *σ* < 0.5 pixels), thus applying the blur in the PSF model to the data. To find the right *σ*, we fitted a subset of the data with sevaral values for *σ* = 0, 0.1,…, 0.5 and selected the *σ* value for which we found neither an accumulation nor a depletion of localizations around the focal plane.

### Post processing

As the positions used above are all based on the objective positions, which differ from the true absolute positions due to refractive index mismatch, we further multiply the *z*-positions with a refractive index mismatch factor of 0.75^1^. Then, *x*, *y*, and *z*-positions were corrected for residual drift by a custom algorithm based on redundant cross-correlation. Localizations persistent in consecutive frames were grouped into one localization, and superresolution images were constructed with every localization rendered as a 2D elliptical Gaussian with a width proportional to the localization precision.

### Sample preparation of clathrin-coated pits in SK-MEL-2 cells

All samples were imaged on round 24 mm high precision glass coverslips No. 1.5H (117640, Marienfeld, Lauda-Königshofen, Germany). Coverslips were cleaned overnight in a 1:1 mixture of concentrated HCl and methanol, rinsed with millipore water until neutral, dried and UV sterilized in a standard cell culture hood.

SK-MEL-2 cells (kind gift from David Drubin, described in Ref. 32) were cultured under adherent conditions in DMEM/F-12 (Dulbecco’s Modified Eagle Medium/Nutrient Mixture F-12) with GlutaMAX and phenol red (ThermoFisher 10565018) supplemented with 10% [v/v] FBS, ZellShield^(tm)^ (Biochrom AG, Berlin, Germany), and 30 mM HEPES at 37°C, 5% CO2 and 100% humidity. Cells were fixed using 3% [w/v] paraformaldehyde (PFA) in cytoskeleton buffer (CB; 10 mM MES pH 6.1, 150 mM NaCl, 5 mM EGTA, 5 mM D-glucose, 5 mM MgCl_2_, described in Ref. 33) for 20 minutes. Fixation was stopped by incubation in 0.1% [w/v] NaBH_4_ for 7 minutes. The sample was washed with PBS three times, and subsequently permeabilized using 0.01% [w/v] digitonin (Sigma-Aldrich, St. Louis, MO, USA) in PBS for 15 minutes. After washing twice with PBS, the sample was blocked with 2% [w/v] BSA in PBS for 60 minutes, washed again with PBS, and stained for 3-12 hours with anti-clathrin light chain (sc-28276, Santa Cruz Biotechnology, Dallas, TX, USA, diluted 1:300) and anti-clathrin heavy chain rabbit polyclonal antibodies (ab21679, Abcam, Cambridge, UK, diluted 1:500) in 1% [w/v] BSA in PBS. The sample was washed with PBS three times, and incubated with a donkey anti-rabbit secondary antibody (711-005-152, Jackson ImmunoResarch, West Grove, PA, USA), which was previously conjugated with Alexa Fluor 647-NHS at an average degree of labeling of 1.5, for 4 hours. Finally, the sample was washed three times with PBS prior to imaging.

For dSTORM imaging, coverslips were mounted in 500 µL blinking buffer (50 mM Tris pH 8, 10 mM NaCl, 10% [w/v] D-glucose, 35 mM 2-mercaptoethylamine (MEA), 500 µg/mL GLOX, 40 µg/mL catalase, 2 mM COT).

### Sample preparation for imaging of the nuclear pore complex and microtubules

Wildtype U-2 OS and genome-edited U-2 OS cells that express Nup107-SNAP (as previously described in Ref. 34) were cultured under adherent conditions in Dulbecco’s Modified Eagle Medium (DMEM, high glucose, w/o phenol red) supplemented with 10% [v/v] FBS, 2 mM L-glutamine, non-essential amino acids, ZellShield^™^ (Biochrom AG, Berlin, Germany) at 37°C, 5% CO_2_ and 100% humidity. All incubations were carried out at room temperature. For nuclear pore staining, the coverslips were rinsed twice with PBS and prefixed with 2.4% [w/v] PFA in PBS for 30 seconds. Cells were permeabilized with 0.4% [v/v] Triton X-100 in PBS for 3 minutes and afterwards fixed with 2.4% [w/v] PFA in PBS for 30 minutes. Subsequently, the fixation reaction was quenched by incubation in 100 mM NH_4_Cl in PBS for 5 minutes. After washing twice with PBS, the samples were blocked with Image-iT^™^ FX Signal Enhancer (ThermoFisher Scientific, Waltham, MA, USA) for 30 minutes. The coverslips were incubated in staining solution (1 µM benzylguanine Alexa Fluor 647 (S9136S, NEB, Ipswich, MA, USA); 1 mM DTT; 1% [w/v] BSA; in PBS) for 50 minutes in the dark. After rinsing three times with PBS and washing three times with PBS for 5 minutes, the sample was mounted for imaging.

For microtubule staining, wildtype U-2 OS cells were prefixed for 2 minutes with 0.3% [v/v] glutaraldehyde in CB + 0.25% [v/v] Triton X-100 and fixed with 2% [v/v] glutaraldehyde in CB for 10 minutes. Fluorescent background was reduced by incubation with 0.1% [w/v] NaBH_4_ in PBS for 7 minutes. After 3 washes with PBS, microtubules were stained using anti alpha-tubulin antibody (MS581, NeoMarkers, Fremont, CA, USA) 1:300 in PBS + 2% [w/v] BSA for 2 h and anti-mouse Alexa Fluor 647 (A21236, Invitrogen, Carlsbad, CA, USA) 1:300 in PBS + 2% [w/v] BSA for 2h. After 3 washes with PBS samples were imaged in a blinking buffer as described above, but with pyranose oxidase instead of glucose oxidase.

For DNA-PAINT imaging, microtubules were labelled with anti alpha-tubulin antibodies (MS581, NeoMarkers, and T6074, Sigma-Aldrich, St. Louis, MO, USA) and anti beta-tubulin antibody (T5293, Sigma-Aldrich) each 1:300 diluted in PBS with 2% [w/v] BSA, for 2 hours. After 3 washes with PBS, samples were incubated with a DNA labelled anti-mouse secondary antibody overnight (docking strand sequence: 5’-TT ATA CAT CTA-3’) and imaged after 5 washes with PBS using 50 pM of complementary Atto-655 labelled DNA imager strand (5’-C TAG ATG TAT-3’-Atto655) in PAINT buffer (PBS, 500 mM NaCl, 40 mM Tris, pH 8.0).

### Embedding of fluorescent beads in agarose gels for calibration measurements

We prepared a 1% [w/v] solution of low melting point agarose (A9414, Sigma-Aldrich) in H_2_O, heated it up to completely dissolve the agarose, and let it cool down to ∼40 °C. We then vigorously vortexed the stock solution of TetraSpeck fluorescent beads (T7279, ThermoFisher Scientific), and added 2.5 µL to 400 µL of the agarose solution, and vortexed again. We then added a 50 µL drop onto a coverslip. After a few minutes, we mounted the sample in H_2_O, and imaged the ∼1 mm thick gel that contained immobilized fluorescent beads throughout.

### Microscopy

SMLM image acquisition was performed at room temperature (24 °C) on a customized microscope^35^ equipped with a high NA oil immersion objective (160x, 1.43-NA oil immersion, Leica, Wetzlar, Germany). We employed a laser combiner (LightHub^®^, Omicron-Laserage Laserprodukte, Dudenhofen, Germany) with Luxx 405, 488 and 638, Cobolt 561 lasers. The lasers were triggered using a FPGA (Mojo, Embedded Micro, Denver, CO, USA) allowing microsecond pulsing control of lasers. After passing through a speckle reducer (LSR-3005-17S-VIS, Optotune, Dietikon, Switzerland), the laser is then guided through a multimode fiber (M105L02S-A, Thorlabs, Newton, NJ, USA). The output of the fiber is first magnified by an achromatic lens and then imaged into the sample^35^. A laser clean-up filter (390/482/563/640 HC Quad, AHF, Tübingen, Germany) is placed in the beam path to remove fiber generated fluorescence. A close-loop focus lock system was implemented using the signal of a near infrared laser reflected by the coverslip and its detection by a quadrant photodiode. The focus can be stabilized within ±10 nm over several hours^36^. The fluorescence emission was filtered by a bandpass filter (700/100, AHF) and recorded by an EMCCD camera (Evolve512D, Photometrics, Tucson, AZ, USA). Typically, we acquire 100,000 – 300,000 frames with 15 ms exposure time (100 ms for DNA-PAINT) and laser power densities of ∼15 kW/cm^2^. The pulse length of the 405 nm laser is automatically adjusted to retain a constant number of localizations per frame.

